# In vivo Tracing of the Ascending Vagal Projections to the Brain with Manganese Enhanced Magnetic Resonance Imaging

**DOI:** 10.1101/2023.01.30.526346

**Authors:** Steven Oleson, Jiayue Cao, Xiaokai Wang, Zhongming Liu

## Abstract

The vagus nerve, the primary neural pathway mediating brain-body interactions, plays an essential role in transmitting bodily signals to the brain. Despite its significance, our understanding of the detailed organization and functionality of vagal afferent projections remains incomplete. In this study, we utilized manganese-enhanced magnetic resonance imaging (MEMRI) as a non-invasive method for tracing vagal nerve projections to the brainstem in vivo and assessing their functional dependence on cervical vagus nerve stimulation (VNS). Manganese chloride solution was injected into the nodose ganglion of rats, and T1-weighted MRI scans were performed 12 and 24 hours post-injection. Our findings reveal that vagal afferent neurons can uptake and transport manganese ions, serving as a surrogate for calcium ions, to the nucleus tractus solitarius (NTS) in the brainstem. In the absence of VNS, we observed significant contrast enhancements of around 19 to 24% in the NTS ipsilateral to the injection side. Application of VNS for four hours further promoted nerve activity, leading to greater contrast enhancements of 40 to 43% in the NTS. These results underline the potential of MEMRI for high-resolution, activity-dependent tracing of vagal afferents, providing a valuable tool for the structural and functional assessment of the vagus nerve and its influence on brain activity.

## Introduction

The vagus nerve, a critical component of the peripheral nervous system, supports rapid communication between the brain and the body’s internal organs. It allows the brain to control physiological functions related to respiratory, cardiovascular, immune, and digestive systems (Chang et al 2015, Powley 2021, Thayer & Lane 2007, Tracey 2002, Travagli & Anselmi 2016). It also allows these organ systems to influence the brain, shaping perception, behavior, cognition, and emotion (Azzalini et al 2019, Craig 2002, Critchley & Harrison 2013, Hsueh et al 2023). Such brain-body interactions are termed as “interoception” (Chen et al 2021), a concept that has recently gained considerable interest in the fields of neuroscience and integrative physiology.

The vagus nerve is being increasingly recognized as a major target of bioelectric medicine (Birmingham et al 2014). Vagus nerve stimulation (VNS), which involves delivering electrical pulses to the vagus nerve at varying levels, has been explored as a therapeutic strategy. It has been used to treat epilepsy (Morris & Mueller 1999), depression (Rush et al 2005), pain (Chakravarthy et al 2015), or to promote learning and rehabilitation (Engineer et al 2019, Engineer et al 2011). It has also gained attention for its potential in relieving chronic conditions affecting internal organs and systems (Bonaz et al 2016, Premchand et al 2014).

However, the mechanism of action for VNS is incompletely understood. The internal structure of the vagus nerve is complex, encompassing nerve fibers with diverse morphological features, fascicular organizations, and functional associations (Havton et al 2021, Stakenborg et al 2020). Optimizing the application of VNS for therapeutic effects often relies on a trial-and-error process. Approximately 80% of vagal nerve fibers are afferent (Berthoud & Neuhuber 2000, Powley et al 2019), enabling sensory neurons in the nodose ganglia (NG) to relay a variety of bodily signals to the nucleus tractus solitarius (NTS) in the brainstem (Prescott & Liberles 2022, Ran et al 2022). These signals, once in the NTS, can further ascend to various brain regions (Berntson & Khalsa 2021, Browning & Travagli 2014, Craig 2002). The interplay between ascending (sensory) and descending (motor) pathways forms the functional neural circuits of interoception, which play a critical role in regulating both mental and physiological states (Chen et al 2021).

Despite the crucial role of the vagus nerve in mediating brain-body interactions, current methodologies constrain our understanding of its structural and functional connectivity. Viral tracing, while useful for localizing neuronal projections (Kaelberer et al 2018), cannot provide functional characterization. Immunohistochemistry with cFos enables the localization of neuronal responses (Cunningham et al 2008), but lacks the capability for longitudinal measures. In vivo electrophysiology offers acute measures of neuronal activity (Cao et al 2021), but is limited by its invasive nature and restricted scope. Functional magnetic resonance imaging (fMRI) offers non-invasive yet indirect measures of neural activity but lacks spatial resolution or specificity to discern fine-grained activations at brainstem nuclei (Cao et al 2017). Diffusion MRI tractography can map structural connectivity but not functional connectivity (Zhang et al 2020). These methodological limitations underscore the need for development and exploration of alternative methods.

Manganese-enhanced magnetic resonance imaging (MEMRI) provides an alternative for potentially addressing the limitations of the aforementioned methods (Pautler et al 1998, Silva et al 2004). The manganese ion (Mn^2+^), being an analogue of the calcium ion (Ca^2+^), can enter the calcium channels of excitable neurons, travel along axonal pathways, enter post-synaptic neurons, and continue to migrate along the circuit (Pautler 2004, Takeda et al 1998). Notably, the transport and uptake of manganese are activity-dependent, providing an index of neural activity (Bearer et al 2007, Duong et al 2000, Lin & Koretsky 1997). Unlike calcium, manganese ions shorten the T_1_ relaxation time, enhancing tissue contrast in T_1_-weighted MRI (Silva et al 2004). Hence, MEMRI has been used to trace active neural pathways in the central nervous system (Chuang et al 2009, Pautler et al 1998, Saleem et al 2002, Watanabe et al 2001, Yu et al 2005). Its application to the peripheral nervous system, however, has been limited. To date, MEMRI has been explored for tracing along the spinal cord and sciatic nerve (Cha et al 2019, Krishnan et al 2020, Matsuda et al 2010), but its utility in tracing the vagus nerve remains largely unexplored.

In the present study, we employ MEMRI to map the ascending vagal pathway, adapting its tracing protocol for the bilateral vagus nerve. We first examined the feasibility of tracing the ascending vagal pathway via the injection of MnCl_2_ (as a source of Mn^2+^) into the left or right nodose ganglion. Next, we verified the activity-dependence of this Mn^2+^ tracing and uptake by modulating vagal activity through cervical VNS. Finally, we quantified the optimal time window for MEMRI. The findings and methodologies presented herein may provide the initial foundation for the in vivo anatomical tracing and functional characterization of the ascending vagal pathway in the rat brain.

## Materials and Methods

### Subjects

This study involved 19 male Sprague-Dawley rats, each weighing between 320 and 400 g. All procedures received approval from our Institutional Animal Care and Use Committee. Prior to surgery, the rats were pair-housed in a single cage within an environment controlled to maintain a 12:12 hour light-to-dark cycle, with the lights turned on at 6 a.m. and turned off at 6 p.m. Following surgery, each rat was housed individually.

### Experiment design

Our study design is demonstrated in Figure 1. The rats were randomly allocated into three groups: group 1 (n=9), group 2 (n=5), and group 3 (n=5). Groups were differentiated based on the tracing of the left vagus nerve (group 1) or the right vagus nerve (group 2) and assessing the effect of the left vagus nerve stimulation (group 3) against a sham control (group 1). All three groups underwent identical surgical and MRI procedures.

**Figure 1.**
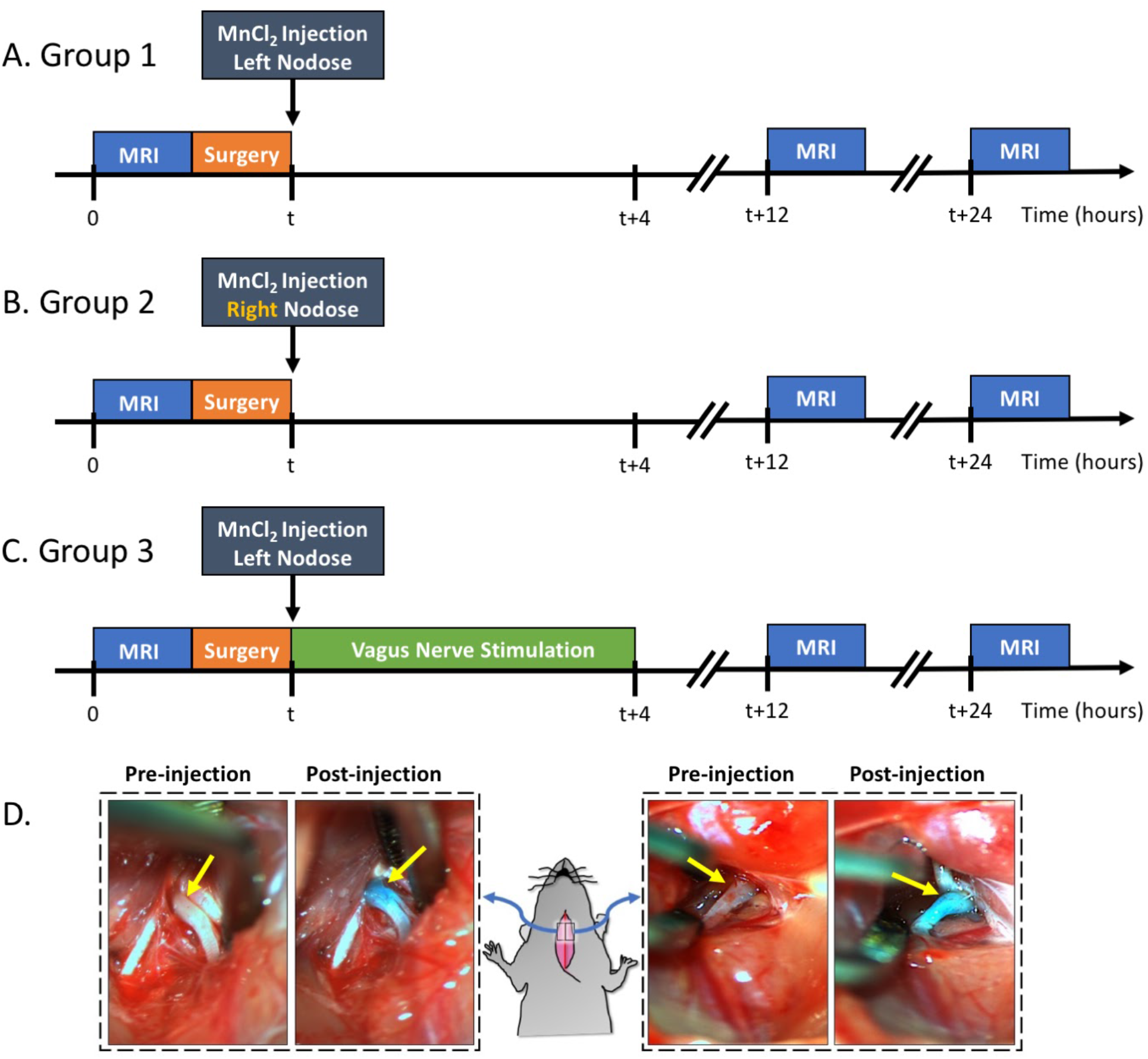
Experiment design and examples of MnCl_2_ injection into NG. A, B, and C illustrate the experimental timelines for groups 1, 2, and 3, respectively. For all three groups, the time of MnCl_2_ injection into either the left or right NG is marked as t. The time of VNS and MRI are marked relative to the time of MnCl_2_ injection. D shows the right and left NG before and after MnCl_2_ injection. The NG appears as green after injection because of the green dye added to the MnCl_2_ solution.

Every rat underwent a T_1_-weighted (T_1_w) brain MRI scan prior to MnCl_2_ injection into the left or right nodose ganglion (NG). Following this pre-contrast MRI, the rats were briefly operated on to expose the NG and the cervical vagus nerve on either the left side (group 1 & 3) or right side (group 2). A solution of MnCl_2_ (1.0 μl of 500 mM, Sigma-Aldrich, St Louis, MO, USA) was injected into the exposed NG. A cuff electrode (Microprobes for Life Science, Gaithersburg, MD, USA) was then wrapped around the exposed vagus nerve. In the case of group 3, electrical stimulation was delivered via the implanted electrode to stimulate the left vagus for 4 hours, after which the electrode was removed. For groups 1 and 2, the electrode was removed without administering any stimulation, thus serving as a sham control for group 3. Post-surgery, all animals were allowed to recover. At 12 and 24 hours post MnCl_2_ injection, each animal underwent another T1w brain MRI scan. The location and level of contrast enhancement due to MnCl_2_ uptake and transport by vagal afferent nerves were assessed by evaluating the voxel-wise intensity difference between pre-contrast and post-contrast MRI.

### Anesthesia, MRI, and surgical protocol

Every rat was initially anesthetized with 5% isoflurane for 5 minutes prior to MRI. The isoflurane was combined with oxygen and delivered at a rate of 500 mL/min. This setting was applicable to other isoflurane dosages mentioned in this protocol unless otherwise stated. During the MRI process, anesthesia was sustained with 1-3% isoflurane. The precise isoflurane dosage varied slightly among animals and was adjusted based on monitored physiological vitals (Model-1030, SA instruments, Stony Brook, NY, USA), such as maintaining the respiratory rate around 40-60 cycles per minute and body temperature at 37-37.5 °C.

For the MRI, each rat was positioned in a customized rat holder in a prone posture with its head secured using a bite bar and two ear bars. Brain MRI was conducted using a 7T horizontal-bore small animal MRI system (BioSpec 70/30; Bruker Instruments, Billerica, USA) fitted with a gradient insert (maximum gradient: 200 mT m^-1^; maximum slew rate: 640 mT^-1^ s^-1^). A ^1^H RF transmit volume coil (86 mm inner diameter) and a ^1^H RF receive-only rat-head surface coil were used for the brain MRI. The imaging session commenced with a localizer to identify the brainstem and other areas of interest. Then a 2D Turbo RARE pulse sequence was used to acquire T_2_-weighted (T2w) images covering the brainstem (TR/TE = 6637.715/32.50ms; FA = 90°; matrix size = 192 × 192; FOV = 32mm × 32mm; slice thickness = 0.438mm; slices = 64; NEX = 2; ETL = 8). A 3D RARE pulse sequence was used to acquire T1w images covering the same region with the same orientation as T2w scans (TR/TE = 300/10ms; FA = 90°; matrix size = 192 × 192 × 64; FOV = 32mm × 32mm × 28mm; NEX = 4; ETL = 8). The same MRI procedure was used both before and after injection of MnCl_2_, generating pre-contrast and post-contrast MRI images for comparison.

Immediately after the pre-contrast MRI, every animal was moved to a surgical station adjacent to the MRI room. Each rat was anesthetized using 5% isoflurane and received an injection of carprofen (10 mg/kg, IP, Zoetis, NJ, USA). The animal was then positioned supine for surgery, with anesthesia maintained at 2% isoflurane. We confirmed adequate anesthesia using a toe pinch reflex test. We then shaved the rat along its neck and cleaned the exposed skin using iodine. An incision was performed along the ventral midline, extending from the mandible to the sternum. The underlying tissue was dissected to expose the trachea and the left or right carotid artery. We identified the cervical vagus nerve next to the carotid artery and further dissected the tissue connecting the carotid artery and the cervical vagus nerve to isolate the vagus nerve. Following the vagus nerve rostrally, we identified the NG.

The NG received a 1.0 μl injection of 500 mM MnCl_2_ solution, administered using a Nanofil 10 μl sub-microliter injection system fitted with a beveled 35-gauge needle (World Precision Instruments, Sarasota, FL). To visually guide and verify the injection, we mixed the MnCl_2_ solution with a green dye (fast green FCF, Sigma-Aldrich, St. Louis, MO, USA). An example of this is shown in Figure 1D. After injection, a bipolar cuff electrode (MicroProbes, Gaithersburg, MD, USA) was wrapped around the exposed vagus nerve.

For the rats in groups 1 and 2, we removed the electrode immediately after implantation and allowed the animals to recover from surgery. However, for those in group 3, we administered vagus nerve stimulation (VNS) for 4 hours. We delivered electrical current pulses via the cuff electrode at an inter-pulse duration (IPD) of 50 ms, a pulse amplitude (PA) of 1 mA, a pulse width (PW) of 0.5 ms, and a frequency of 5 Hz. The stimulation pattern consisted of 20 seconds on and 40 seconds off. After VNS, we removed the electrode, and the animals were allowed to recover. Note that groups 1 and 2 did not receive stimulation but underwent the same surgical procedure and electrode implantation as those in group 3. This made groups 1 and 2 comparable, with group 1 serving as a sham control for group 3.

### Image processing and statistical analysis

For individual animals, we processed MRI data using a combination of FSL (Smith et al 2004), AFNI (Cox 1996), and MATLAB scripts developed in-house. The T_2_-weighted (T2w) images were linearly registered to a rat brain template (Valdes-Hernandez et al 2011) using FLIRT. Furthermore, we registered the T1w images to the T2w images from the same rat using FLIRT. After this, we registered them to the brain template based on the linear transformation determined from the T2w images to the template. After preprocessing, we normalized the intensity of each T1w slice by its average voxel intensity within the same slice.

The voxel-wise intensity increase in the post-contrast T1w MRI was calculated relative to the pre-contrast T1w MRI using the following Equation,

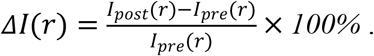

Here, *I*_*post*_ and *I*_*pre*_ represent the voxel intensity at each location *r* in the post- and pre-contrast MRI, respectively. We calculated the relative enhancement Δ*I*(*r*) was calculated for each voxel in each animal and then averaged this across animals in each group.

For each group, we performed a right-tailed paired t-test for comparing the differences between *I*_*post*_ and *I*_*pre*_ for every voxel at a 98% confidence level (or a significance level of alpha = 0.02). The statistically significant voxels were color-coded and shown on the rat brain template to highlight the brain regions where Mn^2+^ ions were taken up.

We also compared the level of enhancement between groups. First, we identified the highest contrast enhancement (in terms of Δ*I*) within a region of interest (primarily the NTS) for each group. We then compared these results between the different groups using a right-tailed two-sample t-test with a confidence level of 95% (alpha = 0.05).

## Results

In our study with rats, we evaluated the potential of MEMRI for tracing vagal afferent projections within the brain. Since vagal afferent neurons are located within the NG, we injected MnCl2 into this region and used MRI to identify and measure T1w contrast enhancement, which results from neuronal uptake and axonal transport of Mn^2+^ ions along the ascending vagal pathways. Figure 2 provides a representative example from a single rat that received a MnCl_2_ injection into its left NG. At the 24-hour mark following the injection, T1w MRI showed notable contrast enhancement at the left NG, the left cervical vagus nerve, and the left NTS, as well as at the right NTS, relative to the pre-injection baseline. The enhancement at the ipsilateral NTS was more substantial than at the contralateral NTS. These results suggest that the Mn^2+^ ions were taken up by sensory neurons in the left NG, transported along the ascending vagal nerves, and then taken up by post-synaptic neurons in the left NTS, which received direct vagal projections. Further uptake by neurons in the right NTS presumably occurred through its neuronal connection with the left NTS.

**Figure 2.**
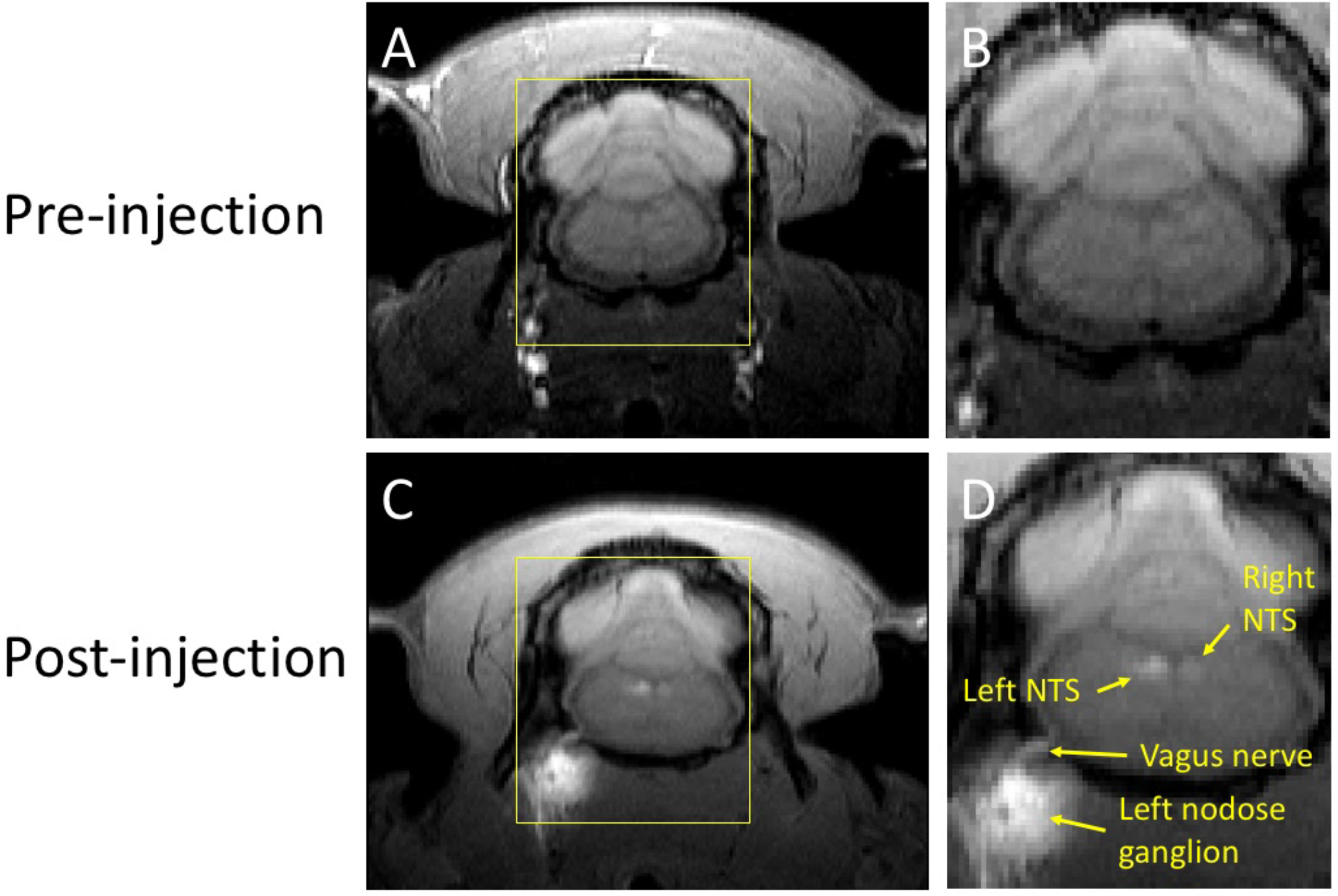
T1w MRI before and after MnCl_2_ injection into the left NG in a single animal. A & C show a single slice of T1w MRI covering the brainstem and NG before and (24 hours) after injection, respectively. B & D zoom in the area within the yellow bounding box. In D, locations of visible enhancement are highlighted with arrows and annotated with anatomical labels.

This finding was corroborated in two animal groups that received MnCl2 injections either in the left (n=9) or the right (n=5) NG. When compared to the pre-injection baseline, the NTS and the area postrema (AP) exhibited significant enhancement in T1w MRI (paired t-test, p<0.02). At the 12-hour mark following the MnCl2 injection, significant contrast enhancement was evident at the ipsilateral NTS and to a lesser degree at the contralateral NTS. At the group level, the strongest contrast enhancement was observed at the ipsilateral NTS, with enhancement levels of 24.75±4.18% and 27.06±1.28% following left and right NG injections, respectively. These findings attest to the high spatial specificity of MEMRI in localizing nuclei that receive vagal afferent projections.

Furthermore, we investigated whether the observed Mn^2+^ uptake was dependent on activity, by applying electrical current pulses (PW=0.5ms; PA=1mA; 5Hz; alternating 20s ON and 40s OFF over a 4-hour period) to the left cervical vagus (group 3, n=5). We then compared the resulting contrast enhancement to the sham group that did not receive stimulation (group 1, n=9). At 12 hours following the MnCl2 injection, the left NTS exhibited the greatest contrast enhancement, with 43.10±6.89% in the presence of VNS relative to 24.75±4.18% without VNS (Figure 4). A similar trend was observed at 24 hours after the MnCl2 injection, where contrast enhancement at the left NTS was higher with VNS (41.74±9.46%) than without VNS (17.28±3.47%). The effect of VNS was significant at both 12 hours (p=0.0160) and 24 hours (p=0.0061) post-MnCl2 injection (Figure 5). These findings suggest that the Mn2+ uptake observed at the NTS with MRI was sensitive to functional changes in vagal nerve activity as a result of VNS.

**Figure 3.**
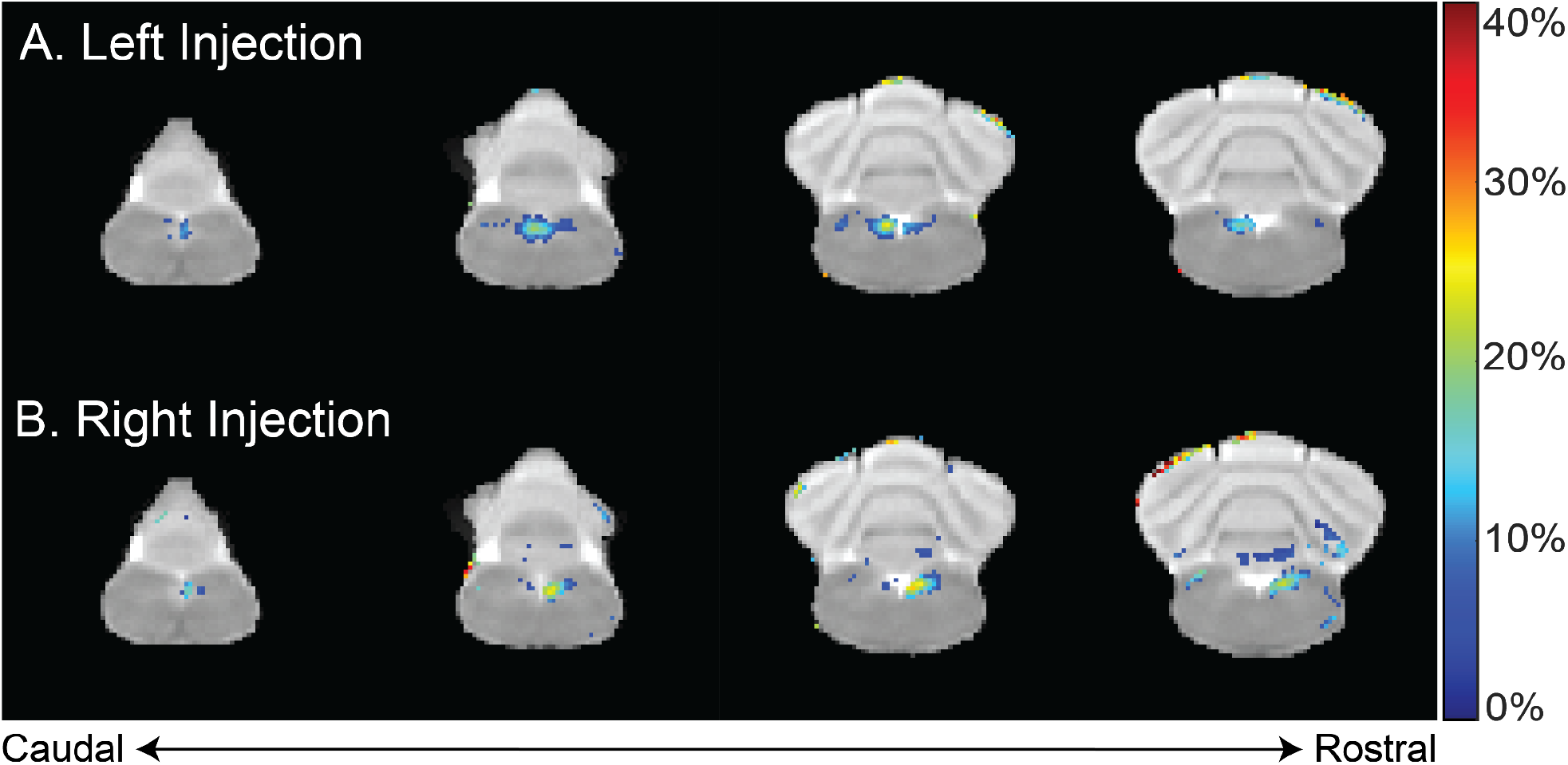
T1w contrast enhancement after MnCl_2_ injection into the left or right NG. A & B show the group-averaged percentage of T1w enhancement at 12 hours after MnCl_2_ injection into the left (group 1) or right (group 2) NG relative to the pre-injection baseline, respectively. Highlighted in color are voxels of statistically significant enhancement (paired t-test, right-tailed p<0.02), where the color indicates the percentage of contrast enhancement. The four slices cover the lower brainstem and cerebellum.

**Figure 4.**
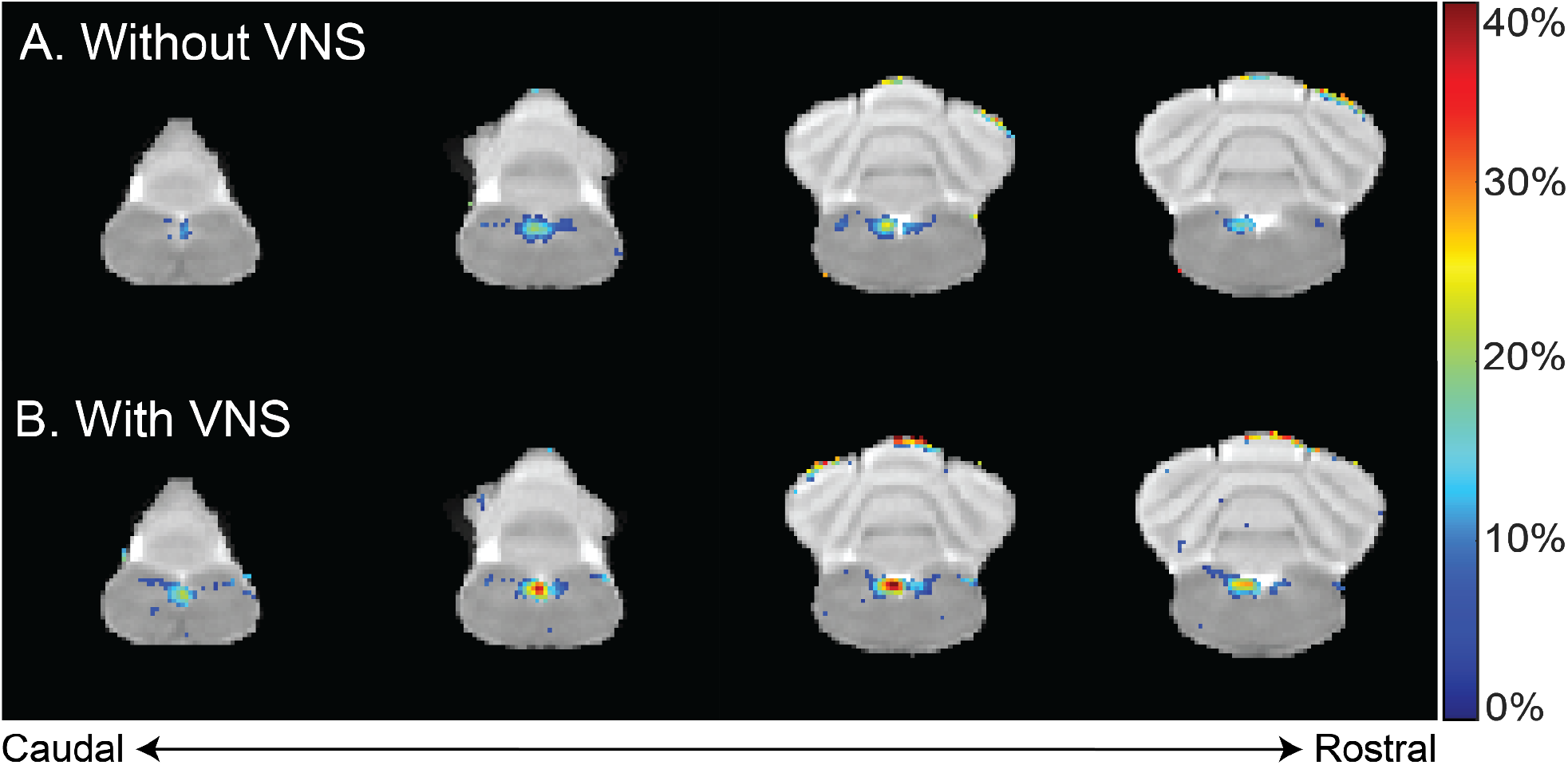
Difference in contrast enhancement with vs. without VNS. A & B summarize the group-averaged contrast enhancement at 12 hours after MnCl_2_ injection into the left NG with or without 4 hours of VNS applied to the left cervical vagus, respectively. Highlighted in color are voxels of statistical significance (paired t-test, right-tailed p<0.02), where the color indicates the percentage of enhancement relative to the pre-injection baseline.

**Figure 5.**
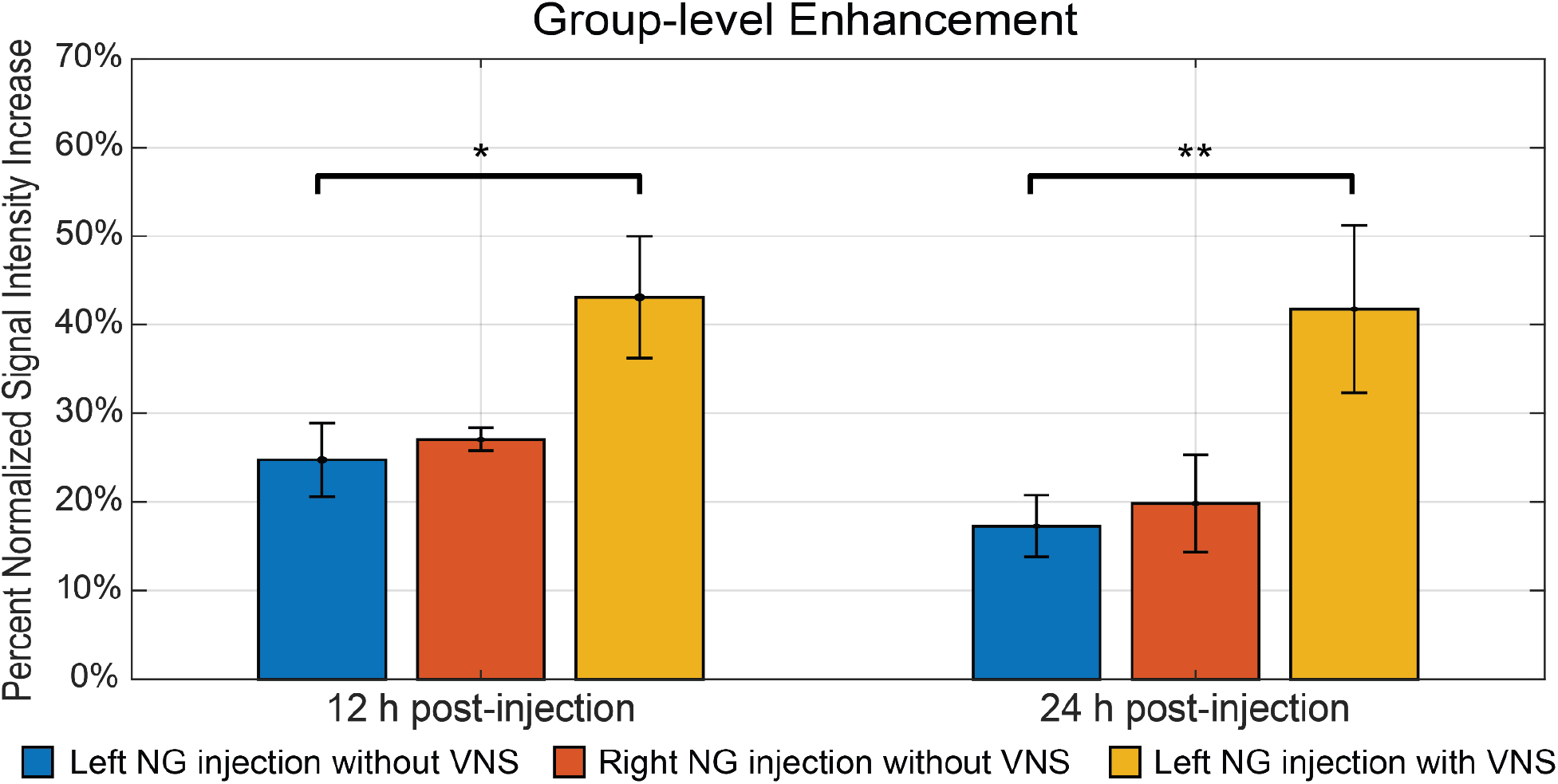
The summary of intensity increase in NTS on the ipsilateral side of nodose ganglion injection. The bar plot shows the mean and standard error of the mean of maximum intensity increase within NTS. The comparison includes all three groups. The white bar indicates group 1, where rats received left nodose ganglion injection but not VNS. The grey bar indicates group 2, where rats received right nodose ganglion injection but not VNS. The black bar indicates group 3, where rats received left nodose ganglion injection and VNS on the same side. The bar plots contain comparisons at both 12 h and 24 h post-injection. * denotes that the difference is statistically significant on the right-tailed two-sample t-test (p<0.05), ** denotes that the difference is statistically significant on the right-tailed two-sample t-test (p<0.01).

Finally, we compared the contrast enhancement at 12 hours and 24 hours post-MnCl_2_ injection to assess the time-dependent uptake, transport, and accumulation of Mn^2+^. Figure 5 depicts the contrast enhancement at the ipsilateral NTS across all three groups. In the absence of VNS, spontaneous vagal activity resulted in slightly higher contrast enhancement at 12 hours post-injection, compared to the 24-hour mark. The effect of VNS was more pronounced at 24 hours, resulting in an additional 24.46% enhancement, compared to 18.35% at 12 hours (Figure 5). This suggests that, following unilateral injection of MnCl_2_ into the NG, Mn^2+^ ions are absorbed by post-synaptic neurons in the bilateral NTS within a 12-hour period, and that MEMRI is highly sensitive to the cumulative effect of vagal afferent activity over a 24-hour period.

## Discussion

In this study, we have demonstrated the utility of MEMRI as an effective technique for mapping vagal afferent projections in rats. By injecting MnCl_2_ into the nodose ganglion (NG) and applying subsequent MRI imaging, we were able to visualize and quantify the T1w contrast enhancement, which signifies the neuronal uptake and axonal transport of Mn^2+^ ions along the vagal nerve pathways. Our findings confirm the robustness and spatial specificity of this technique, showing significant contrast enhancement at the nucleus tractus solitarius (NTS) and area postrema (AP) which receive direct vagal projections. Importantly, we also discovered that Mn^2+^ uptake, as captured by MRI, was sensitive to variations in vagal nerve activity induced by vagus nerve stimulation, underscoring the potential of MEMRI to not only trace neural pathways but also monitor changes in neural activity.

MEMRI provides several unique advantages for studying brain-body interactions mediated by the vagus. Foremost, MEMRI uses MRI and thus is non-invasive. The necessary injection of Mn2+ ions into the nodose ganglion, a peripheral structure housing the cell bodies of vagal afferent neurons, involves a relatively minor surgical intervention without major complications. Post-injection, the animal typically recovers well ahead of the subsequent MRI scans, providing an ample time window (1 to 2 days) for introducing various physiological or pathophysiological states or applying various settings of neuromodulation. The impacts of these states or interventions on vagal nerve projection and activity can be assessed through MEMRI. Moreover, MEMRI can be conducted over multiple sessions, enabling chronic experiments and repeated monitoring of longitudinal changes in neural activity and anatomical connections.

MEMRI presents an opportunity for poly-synaptic neural tracing in an activity-dependent manner, as previous studies tracing CNS pathways have demonstrated (Chuang et al 2009, Pautler et al 1998, Saleem et al 2002, Watanabe et al 2001, Yu et al 2005). In our investigation, we found evidence of Mn^2+^ induced contrast enhancement not only in the ipsilateral NTS, which receives direct vagal projections from the Mn^2+^ injected NG, but also in the contralateral NTS. This observation implies that trans-synaptic uptake of Mn^2+^ occurs by neurons in the ipsilateral NTS. Extensive inter-connections between the left and right NTS further facilitate Mn^2+^ transportation to the opposing side. It is also likely that the left nodose ganglion projects to the right NTS, and vice versa; nevertheless, the contralateral projections are much less common than the ipsilateral projections (Altschuler et al 1989). Also supporting poly-synaptic tracing, we observed contrast enhancement beyond the NTS and AP regions, reaching into other focal areas within the brainstem or cerebellum (data are not shown). However, these additional enhancements lacked consistency across animals and did not achieve statistical significance.

While our current study was unable to consistently trace vagal projections beyond the lower brainstem, we believe that future studies may overcome this limitation through refinement of the contrast agent or the administration method of manganese. A potential strategy could involve the slow release of Mn^2+^ ions, extending the period during which neurons can uptake and transport the manganese injected into the nodose ganglion. Alternative carriers for Mn^2+^ ions such as manganese dipyridoxyl diphosphate (MnDPDP) (Olsen et al 2008, Pochwat et al 2015) or Mn^2+^ encapsulated in nanogels (Eguchi et al 2019) could be explored. Additionally, the use of osmotic pumps could facilitate slow and controlled delivery of Mn^2+^, potentially enhancing the efficacy of tracing (Vousden et al 2018). Such alternatives may also help lower the dose or toxicity of Mn^2+^. With these developments, MEMRI may provide even more comprehensive insights into the dynamic interplay between the vagus nerve and the brain.

## Notes

**Conflict of interest:** The authors declare no competing financial interests.

### Competing Interest Statement

The authors have declared no competing interest.

### Summary of Updates

The manuscript has been updated for better clarification.

## Reference

Altschuler SM, Bao XM, Bieger D, Hopkins DA, Miselis RR. 1989. Viscerotopic representation of the upper alimentary tract in the rat: sensory ganglia and nuclei of the solitary and spinal trigeminal tracts. J Comp Neurol 283: 248–68

Azzalini D, Rebollo I, Tallon-Baudry C. 2019. Visceral Signals Shape Brain Dynamics and Cognition. Trends Cogn Sci 23: 488–509

Bearer EL, Falzone TL, Zhang X, Biris O, Rasin A, Jacobs RE. 2007. Role of neuronal activity and kinesin on tract tracing by manganese-enhanced MRI (MEMRI). Neuroimage 37 Suppl 1: S37–46

Berntson GG, Khalsa SS. 2021. Neural Circuits of Interoception. Trends Neurosci 44: 17–28

Berthoud HR, Neuhuber WL. 2000. Functional and chemical anatomy of the afferent vagal system. Auton Neurosci 85: 1–17

Birmingham K, Gradinaru V, Anikeeva P, Grill WM, Pikov V, et al. 2014. Bioelectronic medicines: a research roadmap. Nat Rev Drug Discov 13: 399–400

Bonaz B, Sinniger V, Hoffmann D, Clarencon D, Mathieu N, et al. 2016. Chronic vagus nerve stimulation in Crohn’s disease: a 6-month follow-up pilot study. Neurogastroenterol Motil 28: 948–53

Browning KN, Travagli RA. 2014. Central nervous system control of gastrointestinal motility and secretion and modulation of gastrointestinal functions. Compr Physiol 4: 1339–68

Cao J, Lu KH, Powley TL, Liu Z. 2017. Vagal nerve stimulation triggers widespread responses and alters large-scale functional connectivity in the rat brain. PLoS One 12: e0189518

Cao J, Wang X, Powley TL, Liu Z. 2021. Gastric neurons in the nucleus tractus solitarius are selective to the orientation of gastric electrical stimulation. J Neural Eng 18: 056066

Cha M, Lee K, Won JS, Lee BH. 2019. Manganese-enhanced magnetic resonance imaging of the spinal cord in rats with formalin-induced pain. Neurosci Res 149: 14–21

Chakravarthy K, Chaudhry H, Williams K, Christo PJ. 2015. Review of the Uses of Vagal Nerve Stimulation in Chronic Pain Management. Curr Pain Headache Rep 19: 54

Chang RB, Strochlic DE, Williams EK, Umans BD, Liberles SD. 2015. Vagal Sensory Neuron Subtypes that Differentially Control Breathing. Cell 161: 622–33

Chen WG, Schloesser D, Arensdorf AM, Simmons JM, Cui CH, et al. 2021. The Emerging Science of Interoception: Sensing, Integrating, Interpreting, and Regulating Signals within the Self. Trends in Neurosciences 44: 3–16

Chuang KH, Lee JH, Silva AC, Belluscio L, Koretsky AP. 2009. Manganese enhanced MRI reveals functional circuitry in response to odorant stimuli. Neuroimage 44: 363–72

Cox RW. 1996. AFNI: software for analysis and visualization of functional magnetic resonance neuroimages. Comput Biomed Res 29: 162–73

Craig AD. 2002. How do you feel? Interoception: the sense of the physiological condition of the body. Nat Rev Neurosci 3: 655–66

Critchley HD, Harrison NA. 2013. Visceral influences on brain and behavior. Neuron 77: 624–38

Cunningham JT, Mifflin SW, Gould GG, Frazer A. 2008. Induction of c-Fos and DeltaFosB immunoreactivity in rat brain by Vagal nerve stimulation. Neuropsychopharmacology 33: 1884–95

Duong TQ, Silva AC, Lee SP, Kim SG. 2000. Functional MRI of calcium-dependent synaptic activity: cross correlation with CBF and BOLD measurements. Magn Reson Med 43: 383–92

Eguchi Y, Murayama S, Kanamoto H, Abe K, Miyagi M, et al. 2019. Minimally invasive manganese-enhanced magnetic resonance imaging for the sciatic nerve tract tracing used intra-articularly administrated dextran-manganese encapsulated nanogels. JOR Spine 2: e1059

Engineer ND, Kimberley TJ, Prudente CN, Dawson J, Tarver WB, Hays SA. 2019. Targeted Vagus Nerve Stimulation for Rehabilitation After Stroke. Front Neurosci 13: 280

Engineer ND, Riley JR, Seale JD, Vrana WA, Shetake JA, et al. 2011. Reversing pathological neural activity using targeted plasticity. Nature 470: 101–4

Havton LA, Biscola NP, Stern E, Mihaylov PV, Kubal CA, et al. 2021. Human organ donor-derived vagus nerve biopsies allow for well-preserved ultrastructure and high-resolution mapping of myelinated and unmyelinated fibers. Sci Rep 11: 23831

Hsueh B, Chen R, Jo Y, Tang D, Raffiee M, et al. 2023. Cardiogenic control of affective behavioural state. Nature

Kaelberer MM, Buchanan KL, Klein ME, Barth BB, Montoya MM, et al. 2018. A gut-brain neural circuit for nutrient sensory transduction. Science 361: eaat5236

Krishnan V, Xu J, Mendoza AG, Koretsky A, Anderson SA, Pelled G. 2020. High-resolution MEMRI characterizes laminar specific ascending and descending spinal cord pathways in rats. J Neurosci Methods 340: 108748

Lin YJ, Koretsky AP. 1997. Manganese ion enhances T1-weighted MRI during brain activation: an approach to direct imaging of brain function. Magn Reson Med 38: 378–88

Matsuda K, Wang HX, Suo C, McCombe D, Horne MK, et al. 2010. Retrograde axonal tracing using manganese enhanced magnetic resonance imaging. Neuroimage 50: 366–74

Morris GL, 3rd, Mueller WM. 1999. Long-term treatment with vagus nerve stimulation in patients with refractory epilepsy. The Vagus Nerve Stimulation Study Group E01-E05. Neurology 53: 1731–5

Olsen O, Thuen M, Berry M, Kovalev V, Petrou M, et al. 2008. Axon tracing in the adult rat optic nerve and tract after intravitreal injection of MnDPDP using a semiautomatic segmentation technique. J Magn Reson Imaging 27: 34–42

Pautler RG. 2004. In vivo, trans-synaptic tract-tracing utilizing manganese-enhanced magnetic resonance imaging (MEMRI). NMR Biomed 17: 595–601

Pautler RG, Silva AC, Koretsky AP. 1998. In vivo neuronal tract tracing using manganese-enhanced magnetic resonance imaging. Magn Reson Med 40: 740–8

Pochwat B, Nowak G, Szewczyk B. 2015. Relationship between Zinc (Zn (2+)) and Glutamate Receptors in the Processes Underlying Neurodegeneration. Neural Plast 2015: 591563

Powley TL. 2021. Brain-gut communication: vagovagal reflexes interconnect the two “brains”. Am J Physiol Gastrointest Liver Physiol 321: G576–G87

Powley TL, Jaffey DM, McAdams J, Baronowsky EA, Black D, et al. 2019. Vagal innervation of the stomach reassessed: brain-gut connectome uses smart terminals. Ann N Y Acad Sci 1454: 14–30

Premchand RK, Sharma K, Mittal S, Monteiro R, Dixit S, et al. 2014. Autonomic regulation therapy via left or right cervical vagus nerve stimulation in patients with chronic heart failure: results of the ANTHEM-HF trial. J Card Fail 20: 808–16

Prescott SL, Liberles SD. 2022. Internal senses of the vagus nerve. Neuron 110: 579–99

Ran C, Boettcher JC, Kaye JA, Gallori CE, Liberles SD. 2022. Publisher Correction: A brainstem map for visceral sensations. Nature 611: E6

Rush AJ, Marangell LB, Sackeim HA, George MS, Brannan SK, et al. 2005. Vagus nerve stimulation for treatment-resistant depression: a randomized, controlled acute phase trial. Biol Psychiatry 58: 347–54

Saleem KS, Pauls JM, Augath M, Trinath T, Prause BA, et al. 2002. Magnetic resonance imaging of neuronal connections in the macaque monkey. Neuron 34: 685–700

Silva AC, Lee JH, Aoki I, Koretsky AP. 2004. Manganese-enhanced magnetic resonance imaging (MEMRI): methodological and practical considerations. NMR Biomed 17: 532–43

Smith SM, Jenkinson M, Woolrich MW, Beckmann CF, Behrens TE, et al. 2004. Advances in functional and structural MR image analysis and implementation as FSL. Neuroimage 23 Suppl 1: S208–19

Stakenborg N, Gomez-Pinilla PJ, Verlinden TJ, Wolthuis AM, D’Hoore A, et al. 2020. Comparison between the cervical and abdominal vagus nerves in mice, pigs, and humans. Neurogastroenterology & Motility 32: e13889

Takeda A, Kodama Y, Ishiwatari S, Okada S. 1998. Manganese transport in the neural circuit of rat CNS. Brain Res Bull 45: 149–52

Thayer JF, Lane RD. 2007. The role of vagal function in the risk for cardiovascular disease and mortality. Biol Psychol 74: 224–42

Tracey KJ. 2002. The inflammatory reflex. Nature 420: 853–9

Travagli RA, Anselmi L. 2016. Vagal neurocircuitry and its influence on gastric motility. Nat Rev Gastroenterol Hepatol 13: 389–401

Valdes-Hernandez PA, Sumiyoshi A, Nonaka H, Haga R, Aubert-Vasquez E, et al. 2011. An in vivo MRI Template Set for Morphometry, Tissue Segmentation, and fMRI Localization in Rats. Front Neuroinform 5: 26

Vousden DA, Cox E, Allemang-Grand R, Laliberte C, Qiu LR, et al. 2018. Continuous manganese delivery via osmotic pumps for manganese-enhanced mouse MRI does not impair spatial learning but leads to skin ulceration. Neuroimage 173: 411–20

Watanabe T, Michaelis T, Frahm J. 2001. Mapping of retinal projections in the living rat using high-resolution 3D gradient-echo MRI with Mn2+-induced contrast. Magn Reson Med 46: 424–9

Yu X, Wadghiri YZ, Sanes DH, Turnbull DH. 2005. In vivo auditory brain mapping in mice with Mn-enhanced MRI. Nat Neurosci 8: 961–8

Zhang Y, Vakhtin AA, Jennings JS, Massaband P, Wintermark M, et al. 2020. Diffusion tensor tractography of brainstem fibers and its application in pain. PLoS One 15: e0213952

